# Rapid Tilt-Series Method for Cryo-Electron Tomography: Characterizing Stage Behavior During FISE Acquisition

**DOI:** 10.1101/2020.05.19.104828

**Authors:** Georges Chreifi, Songye Chen, Grant J. Jensen

## Abstract

We and others recently developed rapid tilt-series acquisition methods for cryo-electron tomography on a Titan Krios G3i equipped with a single axis holder and a K-series direct electron detector and showed that one of these, the fast-incremental single exposure (FISE) method, significantly accelerates tilt-series acquisition when compared to traditional methods while preserving the quality of the images. Here, we characterize the behavior of our single axis holder in detail during a FISE experiment to optimally balance data quality with speed. We explain our methodology in detail so others can characterize their own stages, and conclude with recommendations for projects with different resolution goals.

## INTRODUCTION

Cryo-electron tomography (Cryo-ET) has become an essential imaging technique to reveal the structures of frozen-hydrated biological complexes and the architectures of cells in their native states (Cassidy et al., 2015; Chang et al., 2016; Ghosal et al., 2017; Kaplan et al., 2019; Mattei et al., 2018; Schur et al., 2016). In cryo-ET, a tilt-series is collected by tilting the sample around a tilt axis and acquiring multiple projection images of the specimen throughout a tilt range. This tilt-series is then used to computationally reconstruct a tomogram, a 3D volume of the object being imaged. One of the major challenges in cryo-ET is throughput; a typical tilt-series takes an average of 30 minutes to collect. The majority of this time is spent on ancillary tasks intended to ensure the target remains centered in the field of view (FoV) at every tilt angle in the tilt-series (Mastronarde, 2005). We and others recently developed the fast-incremental single exposure (FISE) method on a Titan Krios G3i equipped with a highly-eucentric single axis holder, and showed that by skipping all tracking of the target during acquisition, high-quality tilt-series can be acquired in 5 minutes or less (Chreifi et al., 2019; Eisenstein et al., 2019). FISE tilt-series of purified ribosomes, for instance yielded subtomogram averages with sub-nanometer resolution (Eisenstein et al., 2019).

During a FISE acquisition, the camera records a single, long movie encompassing the entire tilt-series, including every tilt angle in the tilt scheme, while the dose is fractionated into individual frames at a user-determined framerate. In order to avoid recording images during stage movement, the beam is blanked throughout most of the tilt scheme and only unblanked once the stage has come to a rest at each desired tilt angle. When acquisition is complete, blank frames are discarded based on a user-defined threshold for mean electron count and frames from the same tilt angle are motion-corrected and combined.

While stage movement during tilting has been explored previously (Chreifi et al., 2019; Eisenstein et al., 2019), residual stage movement after the stage has finished tilting to each target angle has not yet been characterized, and could reduce FISE data quality. Here, we have analyzed the settling behavior of our stage immediately after each tilt operation of a typical tilt scheme, with a focus on the direction of movement, its amplitude, how long the settling lasts, and the reproducibility of the behavior for each tilt operation. We find that a fraction of a second delay may be desirable after some tilt operations depending on a user’s data collection parameters and resolution goals. We also describe how we tested our Titan Krios stage at Caltech so others can replicate the tests on their own instruments and customize their FISE scripts accordingly.

## RESULTS & DISCUSSION

In order to fully characterize the behavior of the stage as tilting stops, we first recorded long exposures with the highest frame rate possible starting while the stage was still tilting, and continuing for an additional 2 seconds after the tilting ended (Fig. 1). This experiment was done in triplicates for three different tilt operations commonly used in a typical dose-symmetric tilt scheme: a small 3° tilt operation from 0° to 3°(Fig. 1A) and two large operations in different directions, from 45° to −36° and from −60° to 48° (Fig. 1B,C). In all cases, we observed a quick jolt perpendicular to the tilt axis, with a magnitude of approximately 10 Å, just as the stage appears to arrive at its destination. This jolt was extremely short-lived, lasting for approximately 0.04 s (between 1-3 frames at a frame rate of 75 frames per second (fps)), so it would be easily missed at low frame rates. Afterwards, each of the three tilt operations resulted in different behavior, but in each case, stage movement started at less than 2 Ångstroms per frame and then decreased to hundredths of an Ångstrom per frame within a second.

**Figure 1.**
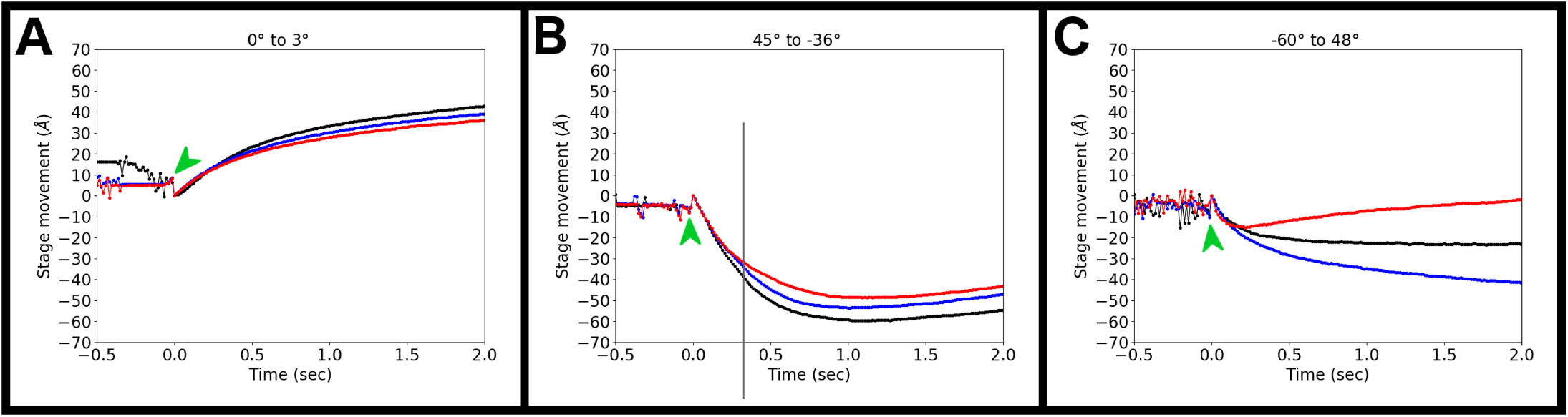
Stage movement perpendicular to the tilt axis during and after tilting from (A) 0° to 3°, (B) 45° to −36°, and (C) −60° to 48°. Time 0 was assigned when the stage appears to have reached its destination. A green arrow marks the jolt observed when the stage stopped tilting.

To further characterize stage stability, we next analyzed residual stage movement in conditions identical to a FISE experiment. In a FISE experiment the serialEM *FrameSeriesFromVar* command is first used to issue a stage tilt command, which can be followed by an optional delay time, and then the beam is unblanked. By setting the delay time to 0 in this experiment, the time between the end of tilting and beam unblanking is set simply by internal communications between the hardware and software. This analysis was done for every tilt operation in a typical −60° to +60°, 3°-increment dose-symmetric tilt scheme (a total of 41 tilt operations, see Table 1 below), and performed at 3 different locations on a grid with cross gratings. For accuracy, the data were again collected at a frame rate of 75 fps, the maximal frame rate achievable by the Gatan K3 camera, and a pixel size of just 0.68 Å/pix. Figure 2 shows residual stage movement after unblanking for the three best and three worst of these 41 tilt operations, in descending order. In the FISE context, the large jolt perpendicular to the tilt axis was never observed in any of the iterations, suggesting that it occurs in the dead time between the end of the stage tilting operation and beam unblanking. The timing of the event therefore suggests that there is no need to account for it in a FISE experiment. In the best cases (Fig. 2A-C, G-I), there was never more than half an Ångstrom movement per frame, even in the first few frames. In the worst cases (Fig. 2D-F, J-L), stage movement began at 2-3 Å/frame and then decreased to about 1 Å/frame after half a second, with further gradual decreases in subsequent seconds. The behavior was very different parallel to the tilt axis. In this direction (Fig. 2M-R, S-X), the stage moved at 0.5-2 Å/frame in the positive direction for the first 2-3 frames (0.1 seconds), but was then as close to motionless as we could detect using standard motion-correction software.

**Table 1.**
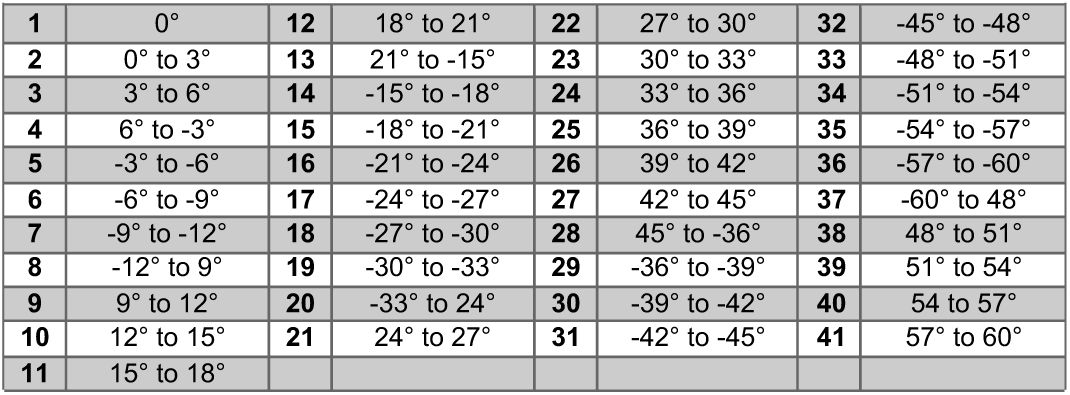
Tilt operations in the dose symmetric tilt-scheme used during FISE acquisition, numbered in the order they are acquired.

**Figure 2.**
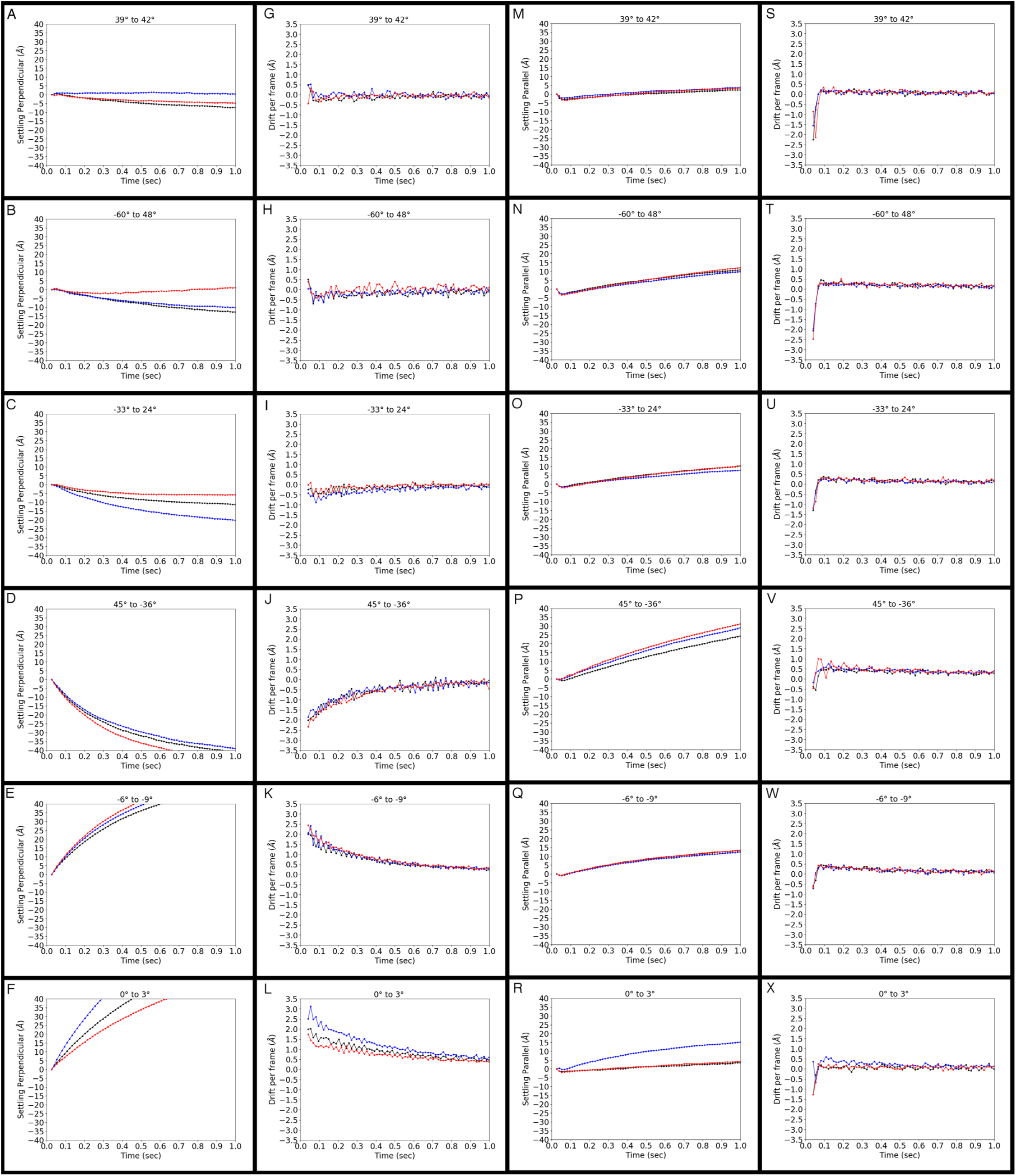
Stage behavior immediately after tilting and unblanking the beam in a FISE experiment for the 3 best and 3 worst tilt operations, in descending order from best to worst. Tilt operations are indicated at the top of each panel. (A-F) Overall stage movement perpendicular to the tilt axis, (G-L) Drift per frame perpendicular to the tilt axis, (M-R) Overall stage movement parallel to the tilt axis, (S-X) Drift per frame parallel to the tilt axis. A small Y-axis scale (± 40 Å) was used in order to show more detailed stage movement during the initial moments following beam unblanking.

Because for the more poorly-performing tilt operations, the stage was still moving after 1 second, we wondered how long one would have to wait for that to stop. To find out, we plotted the movements for 5 seconds and at a larger scale (Fig. 3). The result was that sometimes the stage continued to drift, but not too fast for motion-correction to fix. Thus there is no point to waiting more than just a second or two, since the residual drift after that may persist for a very long time and can be fixed by motion-correction. We were surprised, however, that the worst stage movement perpendicular to the tilt axis followed the smallest tilt operation, the 3° increment from 0° to 3° (Fig. 3F, L). We also thought it remarkable that while the stage’s behavior varied significantly between tilt operations, the movements were quite consistent for a given tilt operation regardless of position on the grid.

**Figure 3.**
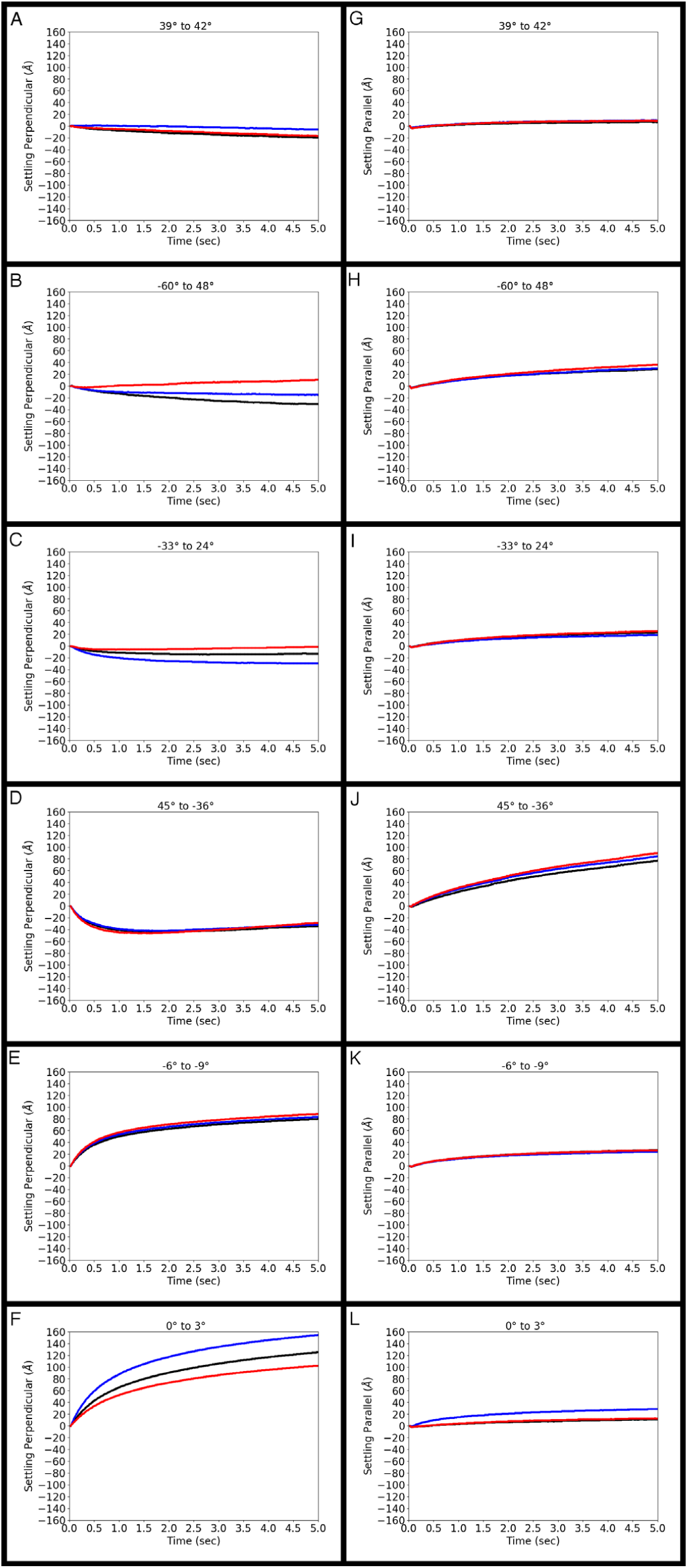
Stage behavior immediately after tilting and unblanking the beam in a FISE experiment for the 3 best and 3 worst tilt operations, in descending order from best to worst. Tilt operations are indicated near the top of each panel. (A-F) Overall stage movement perpendicular to the tilt axis, (G-L) Overall stage movement parallel to the tilt axis.

To examine the quality and resolution of individual frames acquired during a FISE experiment, we generated power spectra at multiple timepoints, ranging from 0.1 s to 5 s, again for the best and worst three tilt operations shown previously (Fig. 2,3). These images were recorded on a cross grating covered with gold nanoparticles. The bright circle is produced by the {111} gold nanoparticle plane (2.35 Å lattice spacing). Another, fainter ring can also sometimes be seen for the {200} plane at 2.03 Å. Two trends in these power spectra were recognized: first, they are missing information perpendicular to the tilt axis at tilt angles above ∼30°, no matter how long the stage is allowed to settle. Because this is the dominant trend, we present the power spectra in Figure 4 in order of final tilt angle reached (rather than as best or worst settling behavior as above). Thus the power spectra for the tilt operations to −36°, 42° and 48° are still incomplete even after 5 seconds (Fig. 4, bottom three rows last column). Information loss perpendicular to the tilt axis at high tilt angles is a well-known phenomenon, though not yet understood in mechanistic detail (Danev et al., 2010; Hagen et al., 2017; Xiong et al., 2009). The second trend visible here, more relevant to the subject of stage movement following tilting, can be seen in power spectra of low-angle tilt operations (top three rows of Fig. 4). In these, frames taken at 0.1 s exhibited only a partial 2.35 Å gold ring, with information missing mostly in the vertical direction, but completeness improves at 0.2 s. Frames taken at 0.5 s and later exhibit complete gold rings at 2.35 Å. These results are in good agreement with the drift data of Figure 2, since drift rates have to decrease to ∼1 Å before resolutions of 2.35 Å can be achieved.

**Figure 4.**
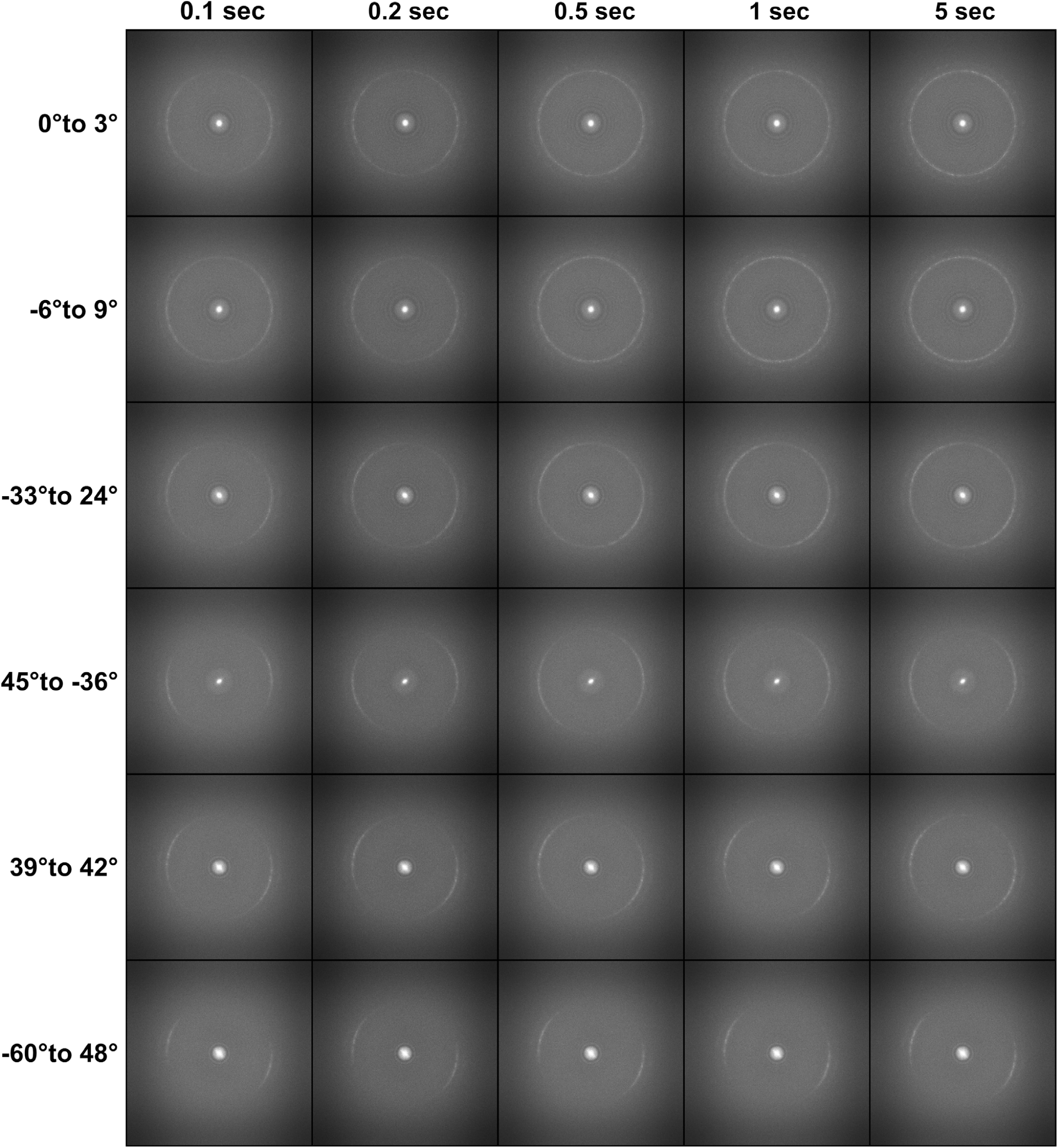
Power spectra of individual frames recorded at 75 fps at various timepoints after beam unblanking for the 3 best and 3 worst tilt operations, shown in order of final tilt angle.

Taken together, these results recommend that short delays be inserted between tilting and unblanking the beam for each specific tilt operation. The ideal delay times are, however, dependent on resolution targets and camera frame rates. This is because one must wait until the absolute drift within a frame falls below about half the resolution target, and this is obviously reached more quickly the faster the frame rate and the larger the resolution target. For every tilt operation described in Table 1, we therefore calculated the time at which residual stage movement between frames drops below certain thresholds, ranging from 1 Å to 5 Å, for frame rates between 10 and 75 fps (Figure 5).

**Figure 5.**
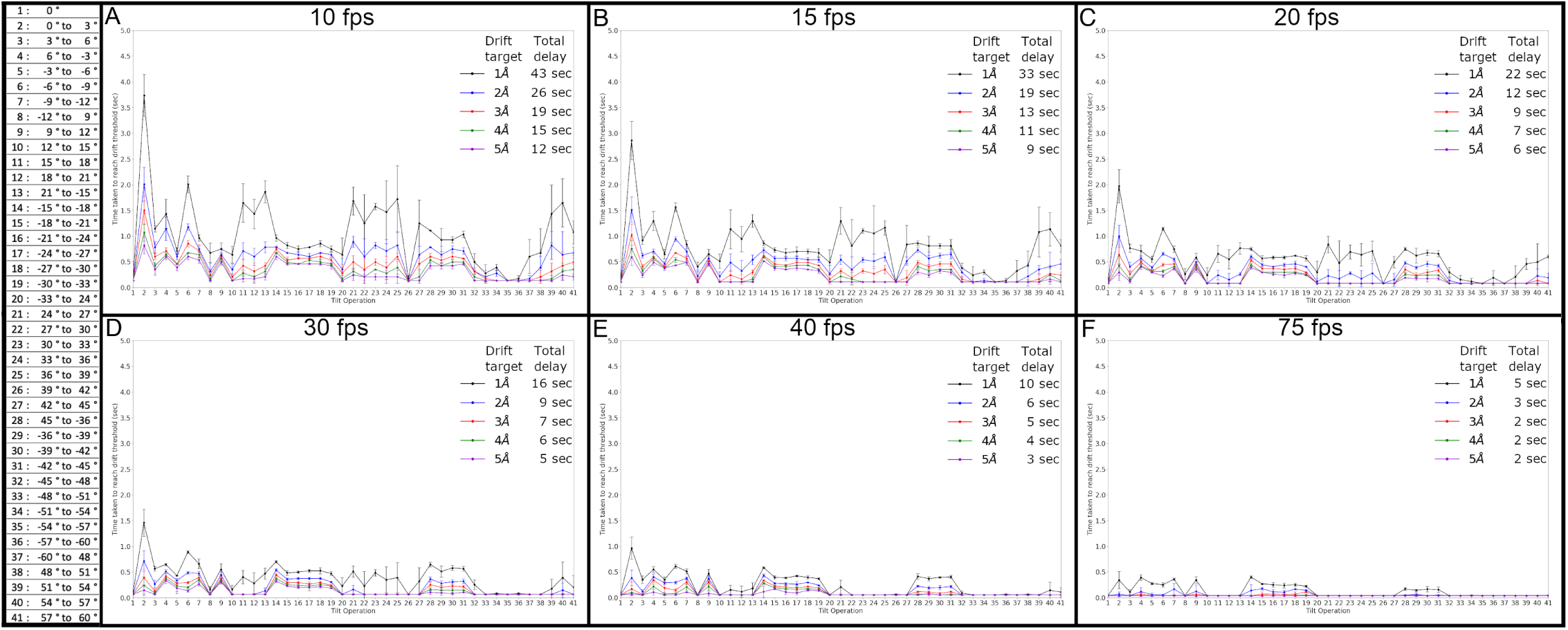
Time required to reach various drift rate thresholds for the 41 tilt operations in a typical dose-symmetric tilt scheme, at frame rates of (A) 10 fps, (B) 15 fps, (C) 20 fps, (D) 30 fps, (E) 40 fps, and (F) 75 fps. For convenience, the attached table references the 41 tilt operations depicted on the horizontal axis of each plot. The inset in each panel shows the color used for each drift rate threshold and the total time that would be added to a FISE tilt-series acquisition if these times were used as delays prior to each tilt operation.

The results highlight how valuable high frame rates can be. Assuming a cryo-ET experiment seeks to produce a sub-nanometer subtomogram average, the drift rate per frame should be less than at least half that resolution and perhaps even as low as a third. Taking a drift rate target of 3 Å/frame, almost no delays are indicated if the camera frame rate is 75 fps (Fig. 5F). If the camera frame rate used is only 10 fps, however, delays of approximately 0.5 seconds are required, adding up to a total of 19 extra seconds needed for the whole tilt-series. Since the achievable resolution in cellular cryo-ET projects is often limited by other factors to just 2-4 nm (Galaz-Montoya and Ludtke, 2017; Glaeser, 1971; Kudryashev et al., 2012; Radermacher, 1988), no delays are needed even at low frame rates.

## CONCLUSIONS

While previous FISE experiments have already generated sub-nanometer resolution subtomogram averages without any delays added (Eisenstein et al., 2019), we suggest an improved FISE protocol that incorporates brief (fractions of a second) delay times between the tilt command and beam unblanking. In order to maximize image quality while minimizing added overall acquisition time, this delay should be tailored to each tilt operation, the resolution goal, and the camera frame rate used. We speculate that different stages will exhibit different behaviors, so each should be characterized following the procedures and analyses we outline here to optimize collection times. We note that modern cryo-stages suffer other shortcomings as well, such as non-eucentricity, which were not discussed here, but can also be corrected by tilt-operation-specific beam and image shifts and focus adjustments in the FISE collection script.

## MATERIALS AND METHODS

### Cryo-ET data collection

All tomographic data were collected in electron-counting mode using *SerialEM* software version 3.8 (Mastronarde, 2003) on a Titan Krios G3i (Thermo Fisher Scientific) equipped with a Gatan energy filter and a K3 electron detector (Gatan). Data were recorded on a Ted Pella waffle pattern grating replica with 2160 lines/mm on 3mm grid (Ted Pella), at a pixel size of 0.68Å and a defocus of −1 µm.

### Data processing and measurement of stage settling

Images were recorded during stage tilting using a simple SerialEM script, by first issuing a *TiltTo* command to each starting tilt angle, followed by a *TiltDuringRecord* command to begin recording while tilting the stage to each target tilt angle, starting from 0° to 3°, 45° to −36°, and −60° to 48°. A long enough exposure time was set in order to record frames during stage tilting and for several seconds after it had stopped. Frames were recorded at a frame rate of 75 fps. Data were then gain-normalized and motion-corrected using IMOD’s *Alignframes* (Kremer et al., 1996). Alignment values produced by the motion-correction were used to plot stage movement perpendicular to the tilt axis as a function of time, and the moment the stage appears to reach the target tilt angle was assigned as time 0.

Data collected in a FISE context were acquired at 75 fps by collecting each tilt angle individually with an exposure time of 5 s per tilt, for a total of 41 datasets at 3 different locations on a waffle pattern grating replica grid (Ted Pella). The dose symmetric tilt scheme used was adapted from others (Hagen et al., 2017), since it provides a more optimal distribution of radiation dose, and is the recommended scheme for FISE data collection. Frames from each dataset were then gain-normalized and motion-corrected with IMOD’s *Alignframes* (Kremer et al., 1996) or Motioncor2 (Zheng et al., 2017). Both motion-correction programs produced similar results in all cases. Motion-correction values were used to plot stage movement perpendicular and parallel to the tilt axis over time. Drift per frame was calculated by taking the difference between the accumulated drift for a given frame and the frame preceding it. Power spectra were produced with EMAN2 using a single 0.0135 s frame at the following timepoints: 0.1 s, 0.2 s, 0.5 s, 1 s, and 5 s. The time taken to reach a drift rate threshold was calculated for each tilt operation by finding the time at which the drift per frame value dropped below a target threshold value and remained below that threshold for at least 10 frames at a given frame rate. Frame rates included 10 fps, 15 fps, 20 fps, 30 fps, 40 fps, and 75 fps, while drift thresholds used were 1 Å, 2Å, 3 Å 4 Å, and 5 Å.

Our FISE scripts can be downloaded at The SerialEM Script Repository (https://serialemscripts.nexperion.net/)

## ACKNOWLEDGEMENTS

This work was supported by NIH grant GM122588 (to G.J.J.). Electron cryomicroscopy was done in the Beckman Institute Resource Center for Transmission Electron Microscopy at Caltech.

